# Quantify the Requirements to Achieve Grain Zn Biofortification of High-yield Wheat on Calcareous Soils

**DOI:** 10.1101/2020.04.09.033639

**Authors:** Sen Wang, Zhaohui Wang, Shasha Li, Chaopeng Diao, Lu Liu, Ning Huang, Ming Huang, Xiaoli Hui, Laichao Luo, Gang He, Hanbing Cao

**Affiliations:** State Key Laboratory of Crop Stress Biology for Arid Areas, Northwest A&F University, Yangling, Shaanxi, China; Key Laboratory of Plant Nutrition and Agri-environment in Northwest China, Ministry of Agriculture and Rural Affairs, College of Natural Resources and Environment, Northwest A&F University, Yangling, Shaanxi, China; Shenzhen Branch, Guangdong Laboratory for Lingnan Modern Agriculture, Genome Analysis Laboratory of the Ministry of Agriculture and Rural Affairs, Agricultural Genomics Institute at Shenzhen, Chinese Academy of Agricultural Sciences, Shenzhen, Guangdong, China

**Keywords:** wheat, grain Zn biofortification, Zn distribution, Zn uptake, quantification

## Abstract

The solution to address global human Zn deficiency is Zn biofortification of staple food crops, aimed at high grain Zn concentration as well as high yield. However, the desired high grain Zn concentration above 40 mg kg^-1^ is rarely observed for high-yield wheat on worldwide calcareous soils, due to inadequate Zn uptake or Zn distribution to grain. The present study aims to investigate how much Zn uptake or distribution is adequate to achieve the Zn.t of high-yield wheat on calcareous soils with low available Zn (∼ 0.5 mg kg^-1^). Of the 123 cultivars tested in a three-year field experiment, 19 high-yield cultivars were identified with similar yields around 7.0 t ha^-1^ and various grain Zn concentrations from 9.3 to 26.7 mg kg^-1^. The adequate Zn distribution to grain was defined from the view of Zn biofortification, as the situation where the Zn distribution to grain (Zn harvest index) increased to the observed maximum of ∼ 91.0% and the Zn concentration of vegetative parts (straw Zn concentration) decreased to the observed minimum of ∼ 1.5 mg kg^-1^ (Zn.m). Under the assumed condition of adequate Zn distribution to grain (∼ 91.0%), all the extra Zn above Zn.m was remobilized from straw to grain and the grain Zn concentration would be increased to its highest attainable level, which was 14.5 ∼ 31.3 mg kg^-1^ for the 19 high-yield cultivars but still lower than 40 mg kg^-1^. Thus, even with the adequate Zn distribution to grain, the current Zn uptake is still not adequate and needs to be increased to 308 g ha^-1^ or higher to achieve Zn.t for high-yield wheat (7.0 t ha^-1^) on low-Zn calcareous soils. Besides, the established method here can also provide the priority measures and quantitative guidelines to achieve Zn biofortification in other wheat production regions.

## 1 Introduction

Although over 50 years have passed since the first case of human zinc (Zn) deficiency (Prasad, 2013), now this problem is still afflicting over 16% of the world population, mostly in Africa and South Asia (Kumssa et al., 2015). In these areas, residents cannot get sufficient Zn intake from daily diets which are mainly low-Zn cereals like wheat, maize, and rice (Wessells and Brown, 2012). Increasing the Zn concentration of cereal grains, known as Zn biofortification, is the main solution to alleviate human Zn deficiency, and the target of Zn fortified wheat grain is 40-50 mg kg^-1^ (Cakmak 2018), much higher than the current worldwide grain Zn concentration of 20-30 mg kg^-1^ (Liu et al., 2014; Chen et al., 2017). To close the gap between the current and target level, it is necessary to fully understand the factors affecting grain Zn concentration.

Grain yield, as an indicator of sink capacity, has essential influence on grain Zn concentration. For instance, 45% decrease in yield of field-grown wheat generated 60% increase in grain Zn concentration (Zhang et al., 2012). The negative correlation between grain Zn concentration and yield has been widely reported in different locations (Liu et al., 2014) and wheat germplasms (Guttieri et al., 2015), and the lowered grain Zn concentration in modern cultivars is mainly caused by the wide adoption of high-yield cultivars after the Green Revolution (Fan et al., 2008; Velu et al., 2017). To feed 9.7 billion population by 2050, the staple crop production has to increase by 60% (FAO, 2014), and thus crop Zn biofortification should increase yield and grain Zn concentration simultaneously (Bouis and Welch, 2010). To achieve high yield and high grain Zn concentration for wheat, it is necessary to enhance both shoot Zn uptake and its distribution to grain (Wang et al., 2018), which are the two sources of Zn accumulated in grain (Kutman et al., 2012; Xue et al., 2012). For example, wheat grain Zn concentration was increased by enhanced nitrogen (N) supply (Kutman et al., 2011; Hui et al., 2017) or ectopic-expressed rice *NICOTIANAMINE SYNTHASE 2* gene (Singh et al., 2017b), because abundant N-containing ligands like nicotianamine (NA) in the phloem enhanced the Zn transportation to grain (Barunawati et al., 2013). Also, the enhanced root Zn uptake increased wheat grain Zn concentration by enlarged root contact area with soil (Ercoli et al., 2017; Singh et al., 2017a), external Zn fertilization (Saha et al., 2017; Liu et al., 2017) or the mobilization of soil intrinsic Zn (Wang et al., 2017). However, most of existing Zn biofortification studies pay much less attention to grain yield, and thus cannot fully achieve the target of Zn biofortification.

The most cost-effective approach to realize crop Zn biofortification is developing cultivars with high yield and high grain Zn concentration (Gregory et al., 2017). For example, a few breeding lines or cultivars of spring wheat exhibited high grain Zn concentration ∼ 40 mg kg^-1^ and high yield ∼ 4 t ha^-1^, under soil available Zn of 1.9 mg kg^-1^ in Canada, India, Pakistan, and Mexico (Gao et al., 2011; Velu et al., 2012). Similarly, under soil available Zn of 1.2 mg kg^-1^ in Iran, several bread wheat genotypes showed high grain Zn concentration > 50 mg kg^-1^ and high yield > 6 t ha^-1^ (Amiri et al., 2015). Thus, under relatively high soil available Zn > 1.0 mg kg^-1^, breeding new wheat cultivars can achieve high grain Zn concentration and high yield simultaneously. But on calcareous soils with low available Zn ∼ 0.5 mg kg^-1^, the success of Zn biofortification of high-yield wheat is rarely achieved. For instance, the highest grain Zn concentration of spring wheat and bread wheat lines was only 25 mg kg^-1^ under soil available Zn of 0.3 ∼ 0.6 mg kg^-1^ in Portugal and Iran (Gomez-Coronado et al., 2016; Khoshgoftarmanesh et al., 2013). In the present study on calcareous soils with available Zn of 0.4 mg kg^-1^ in northwest China, the grain Zn concentrations of 123 wheat cultivars were all below 30 mg kg^-1^. Does this mean that high grain Zn concentration of 40 mg kg^-1^ cannot be achieved by breeding for high-yield wheat on calcareous soils?

The increase of grain Zn concentration for high-yield wheat relies on both high Zn uptake and high Zn distribution to grain (Wang et al., 2018). In the present work, the Zn harvest index varied from 45.5% to 94.0% and the straw Zn concentration at maturity varied from 1.2 to 10.5 mg kg^-1^, while the minimum Zn concentration in wheat vegetative parts was ∼ 5.0 mg kg^-1^ in hydroponics (Kutman et al. 2012), indicating that some Zn in straw might be still transported to grain. Accordingly, inadequate Zn distribution to grain can be the reason for the low grain Zn concentration observed for high-yield wheat. Then, from the view of Zn biofortification, how much Zn distribution to grain is adequate for field-grown high-yield wheat? With adequate Zn distribution, whether the Zn uptake is adequate to achieve high grain Zn concentration (> 40 mg kg^-1^) for high-yield wheat? To test these hypotheses, the Zn concentration, uptake, and distribution of 123 wheat cultivars over three years were analyzed in the present study.

## 2 Materials and methods

### 2.1 Plant materials and field experiment

The field experiment was conducted in Yongshou (108°12′E, 34°44′N) located on the southern Loess Plateau, where the climate is the temperate continental monsoon climate and over half of the annual precipitation (∼ 550 mm) occurs in the summer fallow period from July to September. The top 20 cm soil properties were determined according to Bao (2000) as follows: pH 8.4 (CO_2_-free water), cation exchange capacity 20.3 cmol kg^-1^, soil organic matter 12.9 g kg^-1^, total N 0.9 g kg^-1^, mineral N 26.6 mg kg^-1^, available phosphorus (P) 16.9 mg kg^-1^, available potassium (K) 123.4 mg kg^-1^, available iron (Fe) 7.5 mg kg^-1^, available manganese (Mn) 18.1 mg kg^-1^, available copper (Cu) 1.3 mg kg^-1^, and available Zn 0.4 mg kg^-1^.

One hundred and twenty-three wheat cultivars were collected (Supp Table 1) and tested in the three-year field experiment with a randomized complete block design. Each cultivar had four replications and each plot consisted of four 200 cm long rows with 2.5 cm seed spacing and 20 cm row spacing, under the nutrient supply of 150 kg N ha^-1^ and 100 kg P_2_O_5_ ha^-1^. All cultivars were sown during 28th to 30th September in 2013, 2nd to 3rd October in 2014, and 26th to 28th September in 2015, and harvested during 14th to 17th June in 2014, 18th to 20th June in 2015, and 18th to 20th June in 2016. The rainfalls during summer fallow periods and growing seasons were 271 and 267 mm in 2013-2014, 317 and 314 mm in 2014-2015, and 228 and 186 mm in 2015-2016, respectively. Throughout the experimental period, no irrigation was conducted, and herbicide and pesticide were used when necessary.

### 2.2 Sampling and chemical analyses

At maturity, the plants of 30 ears were randomly sampled from the central two rows of each cultivar plot, and the roots were cut off at the stem-root joint part. Then, the shoots were air-dried and separated into stems, glumes, and grains, which were washed with deionized water and oven-dried at 65°C to determine dry weight. After that, the oven-dried samples were ground by a ball mill (Retsch MM400, Germany) for chemical analyses. The remaining ears in the central two rows of each plot were also harvested and weighted, plus the above grain weight, to estimate the grain yield, glume biomass, and stem biomass in oven-dried weight.

Plant Zn concentration was determined as Ozbek and Akman (2016) with some modifications. Briefly, 0.2 g sample was mixed with 5.0 ml HNO_3_ (65%) in a 50 ml Teflon tube and predigested at 120°C for 0.5 h, and then 1.0 ml H_2_O_2_ (30%) was added before microwave digestion (MW Pro, Anton Paar, Austria). Each sample was digested in two technical duplicates and the standard wheat flour (GBW10011 GSB-2) was used for quality control. The digestion solution was diluted with ultrapure water (18.25 MΩ cm^-1^), and Zn concentration was determined by ICP-MS (iCAP Qc, Thermo Fisher Scientific, USA). The measured Zn isotope was ^66^Zn, and ^73^Ge was added as internal standard to calibrate the signal fluctuation.

### 2.3 Quantification of the potential increase of grain Zn concentration by adequate Zn distribution

From the perspective of Zn biofortification, the adequate Zn distribution to grain refers to the situation where nearly all the mobilizable Zn in shoot vegetative parts was transported or remobilized to grain, and correspondingly, the ratio of grain Zn uptake to shoot Zn uptake (Zn harvest index) increased to the maximum and the Zn concentration of vegetative parts (straw Zn concentration, Zn.s) decreased to the minimum. To determine the adequate Zn distribution to grain for high-yield wheat, we selected the cultivars with yields higher than the corresponding median of all cultivars in each year, calculated the Zn uptake and Zn harvest index, and combined stem and glume Zn concentrations into Zn.s by the weighted mean method. In each year, the observed maximum Zn harvest index or minimum straw Zn concentration (Zn.m) was used to indicate the level of adequate Zn distribution to grain. Since Zn concentration was easier to measure than Zn harvest index, we used Zn.m to calculate the potential increase of grain Zn concentration (Zn.p) caused by the extra Zn remobilized from straw to grain under the condition of adequate Zn distribution to grain.

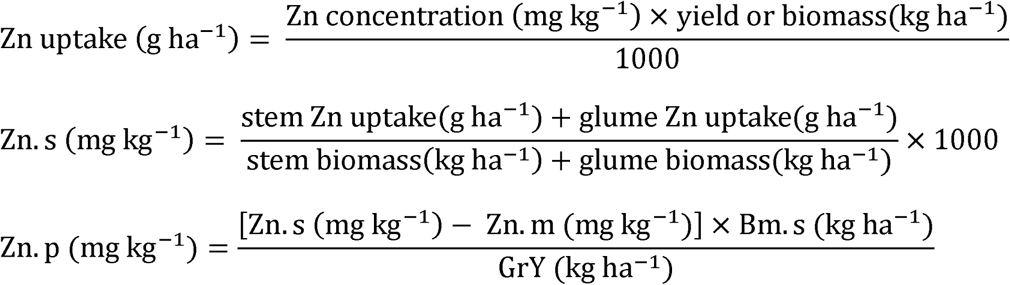

Here Bm.s and GrY refer to straw biomass and grain yield. The sum of Zn.p and the current grain Zn concentration (Zn.c) was defined as the attainable grain Zn concentration (Zn.a).

Shoot Zn uptake and Zn harvest index (ZnHI) were calculated as described by Kutman et al. (2011) and Xue et al. (2012).

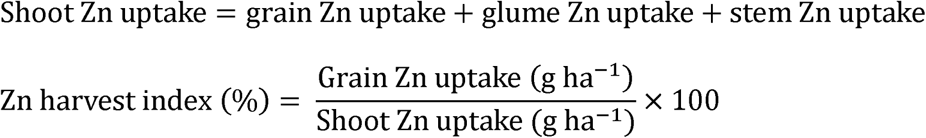

### 2.4 Statistical analyses and graphing

Analysis of variance (ANOVA) was used to test the effects of year, cultivar, and year × cultivar interaction on yield or biomass, Zn concentrations, and Zn uptakes. The relationships of grain Zn concentration with other traits were tested by Pearson correlation or linear regression. All the statistical analyses were completed by PROC GLM, PROC CORR or PROC REG in SAS v 8.01, with 0.05 set as the significance level. Scatter and bar plots were created by Sigmaplot 12.5, and the concept chart illustrating Zn biofortification guidelines was created in Adobe Illustrator CC.

## 3 Results

### 3.1 Relationship of grain Zn concentration with other traits for high-yield wheat

Of the 123 tested wheat cultivars ordered by yield from low to high, 19 cultivars consistently exhibited higher yields than the median in each year, and were identified as high-yield cultivars (Figure 1). Although yield and biomasses showed significant differences among three years, the 19 high-yield cultivars exhibited similar yields (∼ 7 t ha^-1^) and varied greatly in grain Zn concentration, from 14.7 to 26.7 mg kg^-1^ in 2014, 9.3 to 22.2 mg kg^-1^ in 2015, and 13.5 to 17.0 mg kg^-1^ in 2016 (Supp Table 2 and Figure 2A). Besides, grain Zn concentration showed no relation with yield, the biomass and Zn concentration of stem and glume (Figure 2A and Supp Figure 1). However, shoot Zn uptake was positively correlated with grain Zn concentration in each year and Zn harvest index was positively correlated with grain Zn concentration in 2015 (Figure 2C and 2D).

**Figure 1.**
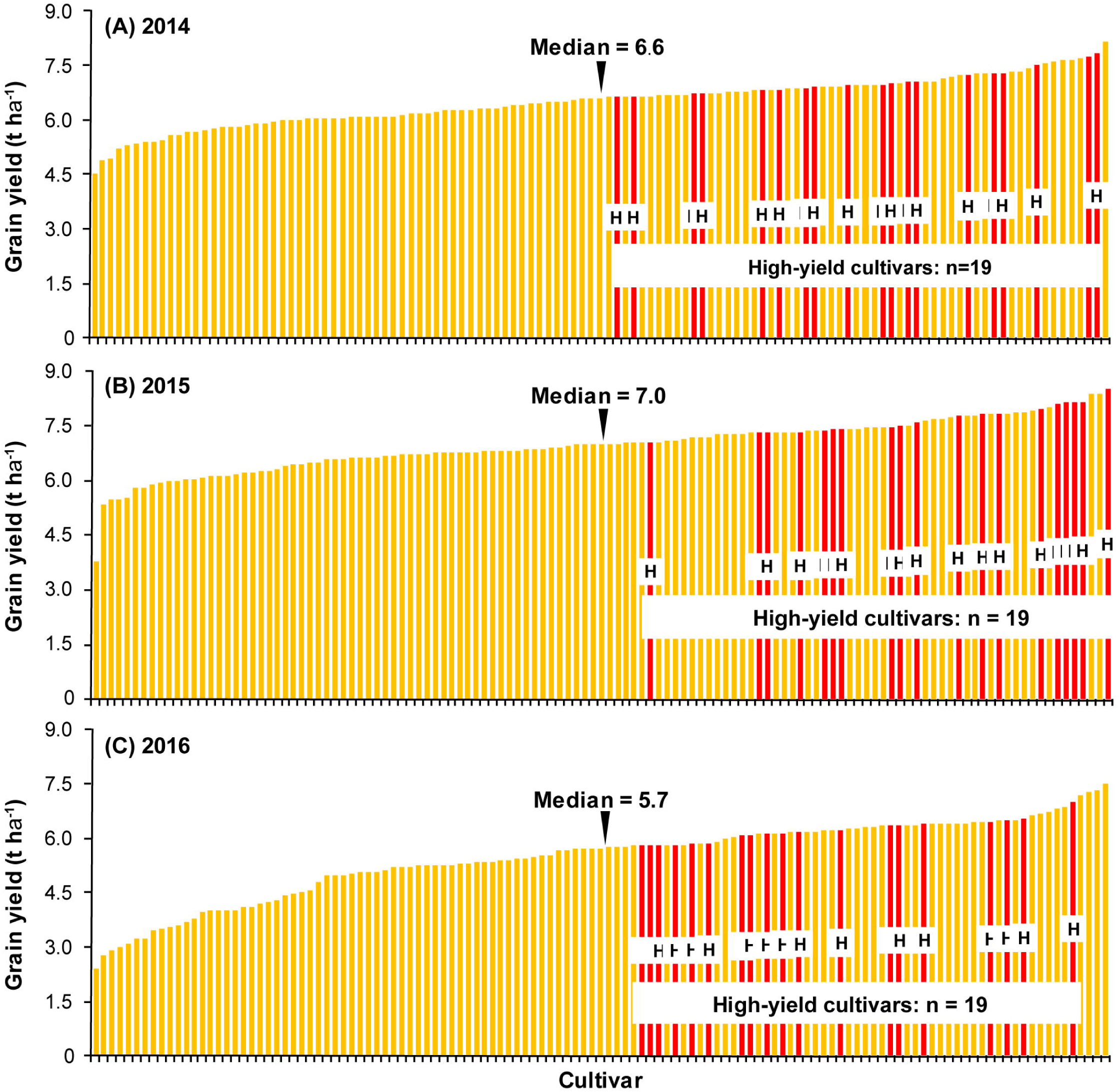
Yield variations of the 123 tested wheat cultivars in 2014 (A), 2015 (B), and 2016 (C). The 19 high-yield cultivars (red bar, ‘H’) were identified with their yields consistently higher than the corresponding median of all cultivars in each year.

**Figure 2.**
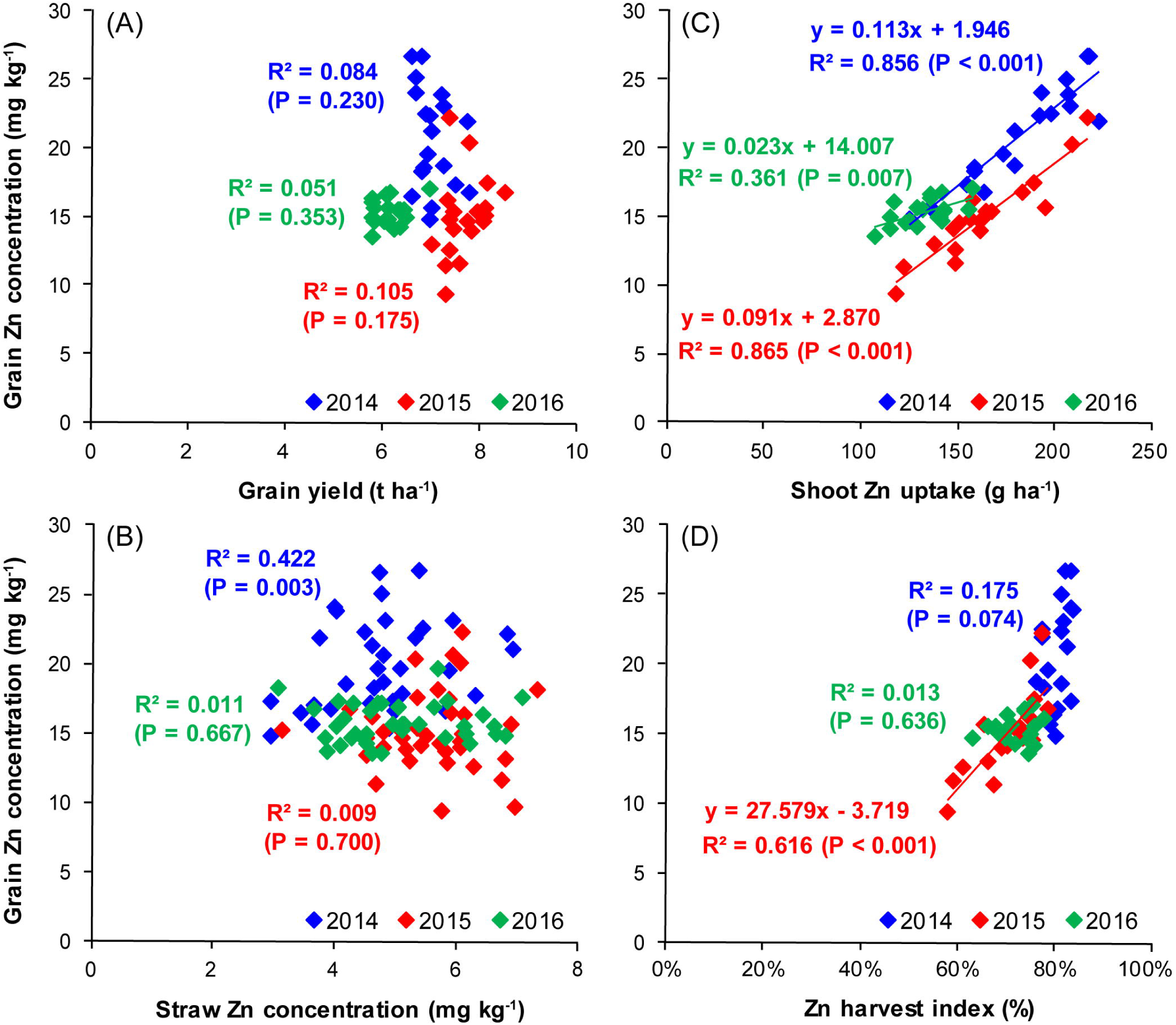
Relationships of grain Zn concentration to grain yield (A), straw Zn concentration (C), shoot Zn uptake (B), and Zn harvest index (D) for the high-yield wheat cultivars over three experimental years. Determination coefficients (*P*-values) indicate the relationships of grain Zn concentration to these traits, and the significant regressions at *P* < 0.05 are shown by solid lines.

### 3.2 Minimum straw Zn concentration and maximum Zn harvest index

Shoot Zn uptake and Zn harvest index showed large variations among the 19 high-yield cultivars or three experimental years (Supp Table 2, Figure 3A and 3B). The observed maximum Zn harvest index was 91.8% in 2014, 87.7% in 2015, and 94.0% in 2016 (Figure 3B), while the observed minimum straw Zn concentration (Zn.m) was 1.3 mg kg^-1^ in 2014, 1.9 mg kg^-1^ in 2015, and 1.2 mg kg^-1^ in 2016 (Figure 3C). With Zn.m used to indicate the level of adequate Zn distribution to grain, the gap between current straw Zn concentration and Zn.m was used to calculate the amount of extra Zn which could be remobilized into grain.

**Figure 3.**
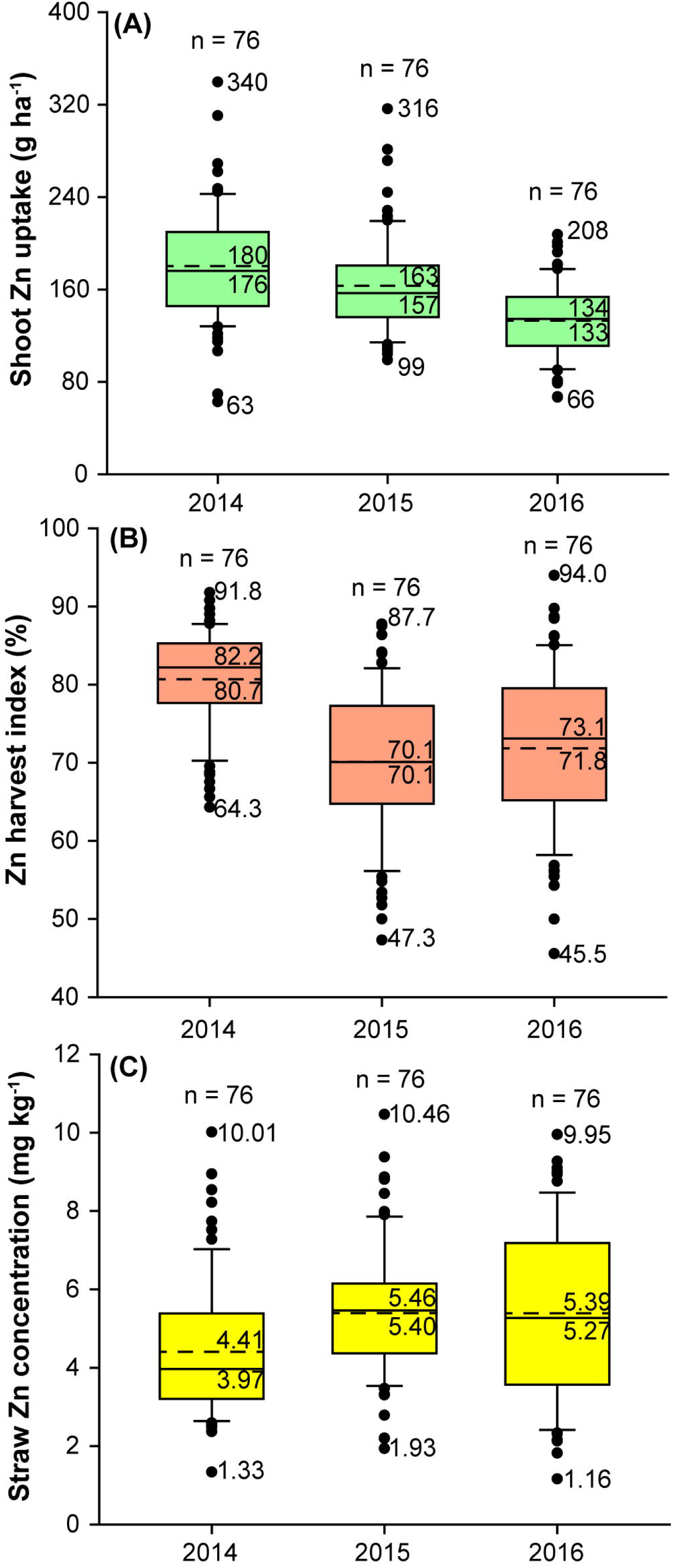
Variations of shoot Zn uptake (A), Zn harvest index (B), and straw Zn concentration (C) of the high-yield cultivars over three experimental years. For each box plot, like the straw Zn concentration in 2014, the five values from top to bottom are the number of original observations (19 cultivars × four replications = 76), the maximum (10.01), the median (4.41, dashed line), the average (3.97, solid line) and the minimum (1.33), respectively.

### 3.3 Potential increase of grain Zn concentration and attainable grain Zn concentration

The potential increase of grain Zn concentration (Zn.p) due to extra Zn remobilized from straw to grain of the 19 high-yield cultivars ranged from 2.1 to 5.5 mg kg_-1_ in 2014, 3.0 to 6.5 mg kg^-1^ in 2015, and 3.5 to 7.5 mg kg^-1^ in 2016 (Figure 4). When Zn.p was added to the current grain Zn concentration (Zn.c), the attainable grain Zn concentration (Zn.a) varied from 17.0 to 31.3 mg kg^-1^ in 2014, 14.5 to 27.5 mg kg^-1^ in 2015, and 17.6 to 22.8 mg kg^-1^ in 2016 (Figure 4), lower than the Zn biofortification target (Zn.t) of 40 mg kg^-1^. Thus, even with the adequate Zn distribution to grain, the current shoot Zn uptake was not adequate to ahieve Zn.t of high-yield wheat. For an assumed high-yield (∼ 7 t ha^-1^) wheat cultivar with adequate Zn distribution to grain (∼ 91.0%), the required shoot Zn uptake should be at least 308 g ha^-1^ to achieve the Zn.t.

**Figure 4.**
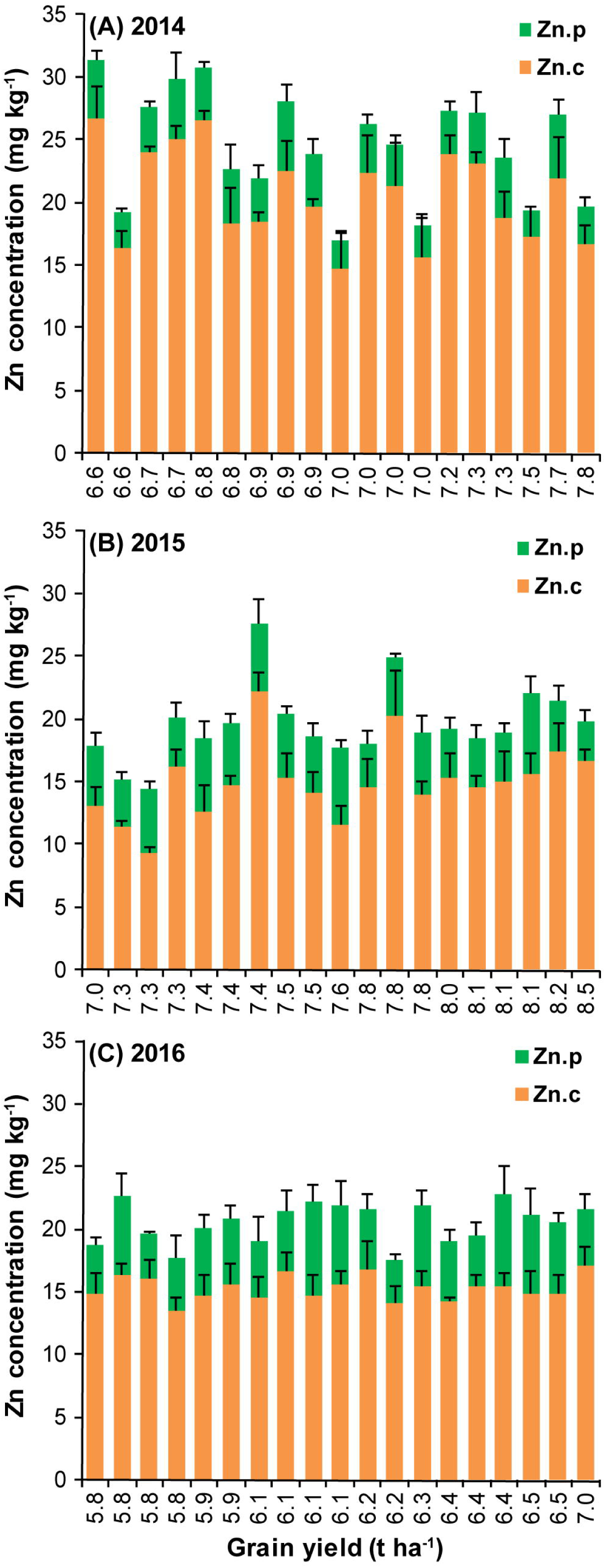
Grain Zn concentration (Zn.c, orange bar) and its biofortification potential (Zn.p, green bar) of the 19 high-yield wheat cultivars in 2014 (A), 2015 (B), and 2016 (C). Error bars stand for standard error (n=4).

## 4 Discussion

### 4.1 Using the minimum straw Zn concentration to estimate the adequate Zn distribution to grain

In the present work, the adequate Zn distribution to grain is defined from the view of Zn biofortification, and it refers to the situation where all the mobilizable Zn in shoot vegetative parts was transported or remobilized to grain, meaning that Zn harvest index increased to its maximum and straw Zn concentration decreased to its minimum (Zn.m). Then, the gap between current straw Zn concentration and Zn.m can be used to calculate the amount of extra remobilized Zn from straw to grain, which results in the potential increase and the attainable highest grain Zn concentration under specific conditions. For high-yield bread wheat grown on low-Zn calcareous soils, the observed minimum straw Zn concentration is used as Zn.m which is ∼ 1.5 mg kg^-1^ and lower than the reported 5.0 mg kg^-1^ for durum wheat in hydroponics (Kutman et al., 2012). Presumably, the senescent vegetative parts of field-grown wheat need less Zn to constitute structural components and more Zn can be remobilized into grain. Under uniform environmental conditions, straw Zn concentration and Zn.m can be used to determine whether the Zn distribution to grain is adequate or not to promote Zn biofortification. Since the Zn biofortification of high-yield wheat needs both high Zn uptake and high Zn distribution to grain (Wang et al., 2018), the Zn concentration of vegetative parts, which is easier to measure than the Zn harvest index, deserves to be considered in future studies on wheat Zn biofortification.

### 4.2 Quantify the requirements to achieve wheat Zn biofortification in different environments

Based on the adequate Zn distribution to grain (∼ 91.0%) as indicated by Zn.m, we can get the attainable highest grain Zn concentration (Zn.a) and compare it with the grain Zn biofortification target (Zn.t, 40 mg kg^-1^), which can tell us whether the current Zn uptake is adequate to achieve Zn.t for high-yield wheat (∼ 7 t ha^-1^). Besides, the established quantitative analysis method here are also suitable for other wheat production regions with the aim of increasing the current grain Zn concentration (Zn.c) and grain yield. The priority measures to achieve the Zn.t of high-yield wheat are different under different conditions (Figure 5). In wheat planting areas where Zn.a is close to or higher than Zn.t and the soil available Zn is often relatively high (> 1.0 mg kg^-1^), the priority to realize Zn.t is to adopt or develop the cultivars with high Zn distribution to grain. For example, in North China Plains with soil available Zn of 3.4 mg kg^-1^, optimal N application generated high yield of 7.5 t ha^-1^ and shoot Zn uptake of 370 g ha^-1^ (Xue et al., 2014), and the Zn.a would be 44.9 mg kg^-1^ and higher than Zn.t, indicating the necessity to introduce high-Zn-distribution wheat cultivars. In wheat production regions where Zn.a is lower than Zn.t and higher than Zn.c and the soil available Zn is often relatively low (< 0.5 mg kg^-1^), developing high-Zn-distribution cultivars should be combined with agronomic measures like Zn fertilization to promote Zn uptake. For instance, in Iran rainfed drylands with soil available Zn of 0.2 mg kg^-1^ (Norouzi et al., 2014), the tested wheat cultivars exhibited high yield of 6.6 t ha^-1^ and shoot Zn uptake of 174 g ha^-1^. Even if the Zn.c of 17.0 mg kg^-1^ was increased to the Zn.a of 26.4 mg kg^-1^ by introducing new cultivars, there was still a large gap to Zn.t which needed to be closed by increasing Zn uptake. In wheat production regions where Zn.a is close to Zn.c and lower than Zn.t, the Zn distribution to grain has reached its maximum and the priority measure of Zn biofortification is to increase Zn uptake by agronomic practices. However, this case is rarely observed up to now because researchers have not paid enough attention to the Zn in vegetative parts in wheat Zn biofortification studies.

**Figure 5.**
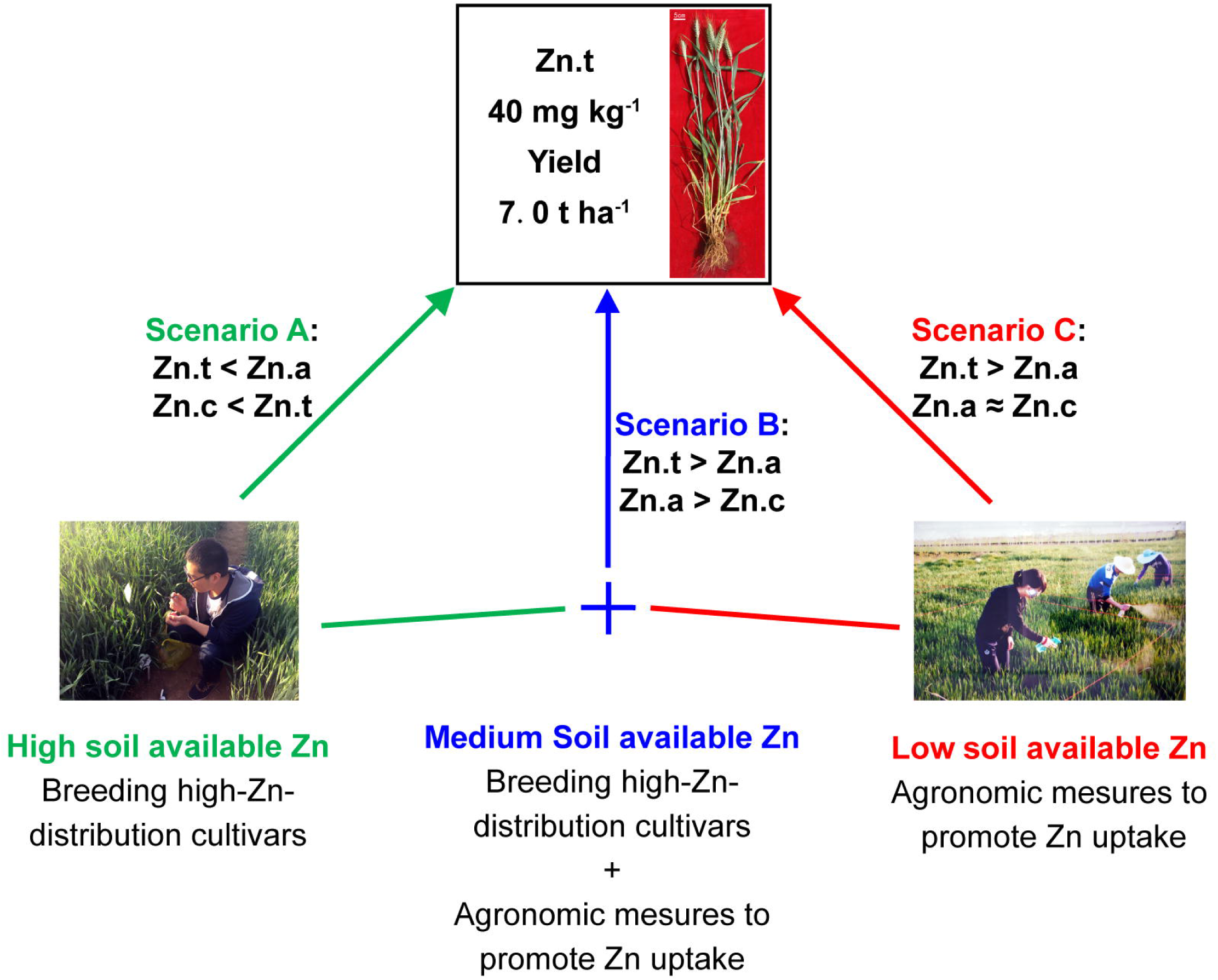
The most suitable Zn biofortification measures in different scenarios of the current grain Zn concentration (Zn.c), attainable grain Zn concentration (Zn.a), and Zn biofortification target level (Zn.t). This is proposed as a guiding workflow to realize grain Zn biofortification target for high-yield wheat.

On calcareous soils with low available Zn < 0.5 mg kg^-1^, the current wheat grain Zn concentration is around 20 mg kg^-1^, as reported in China, Kazakhstan, Mexico, Turkey, and Zambia, etc. (Zou et al., 2012; Liu et al., 2014). In the present study on calcareous soils with available Zn of 0.4 mg kg^-1^, the wheat cultivar Zhengmai 7698 had the highest Zn.a of 31.3 mg kg^-1^ and the yield of 6.6 t ha^-1^ in 2014, which was still lower than the Zn.t. To achieve high grain Zn concentration of 40 mg kg^-1^ and high yield of 7.0 t ha^-1^ simultaneously, the current shoot Zn uptake should be increased to at least 308 g ha^-1^ by agronomic measures like soil Zn fertilization with the wheat cultivars with high Zn distribution to grain (∼ 91%). These guidelines deserve to be tested in field studies on Zn biofortification.

## 5 Conclusion

From the view of crop Zn biofortification, the adequate Zn distribution to grain can be defined as the case where Zn harvest index increased to its maximum and straw Zn concentration decreased to its minimum. For the high-yield (∼ 7 t ha^-1^) wheat grown on low-Zn (∼ 0.5 mg kg^-1^) calcareous soils, the maximum Zn harvest index was ∼ 91.0% and the minimum straw Zn concentration was ∼ 1.5 mg kg^-1^. Under the condition of adequate Zn distribution to grain, the gap between straw Zn concentration and its minimum could be used to determine the extra Zn remobilized to grain and the highest attainable grain Zn concentration, which was 14.5 ∼ 31.3 mg kg^-1^ for the high-yield wheat cultivars and lower than the grain Zn biofortification target (Zn.t) of 40 mg kg^-1^. Therefore, even with the adequate Zn distribution to grain (∼ 91%), the current shoot Zn uptake is still not adequate to achieve the Zn.t of high-yield wheat grown on low-Zn calcareous soil, and the priority measure of Zn biofortification is to increase Zn uptake to 308 g ha^-1^ or higher by agronomic practices like Zn fertilization. The established method here can also provide the most suitable guidelines and quantitative requirements to achieve grain Zn biofortification in other wheat production regions.

## 6 Abbreviations

Zn.s: straw Zn concentration
Zn.m: minimum straw Zn concentration
Zn.p: potential increase of grain Zn concentration
Zn.c: current grain Zn concentration
Zn.a: attainable grain Zn concentration
ZnHI: Zn harvest index
Zn.t: grain Zn biofortification target.

## 8 Author Contributions

SW, ZHW, SSL, CPD, and LL contributed conception and design of the study. SW, SSL, CPD, LL, NH, MH, XLH, LCL, GH, and HBC conducted experiment, sampling, chemical and statistical analyses. SW wrote the first draft of the manuscript and ZHW, MH, XLH, LCL, GH, and HBC read and revised the manuscript. All authors contributed to manuscript revision, read and approved the submitted version.

## 10 Acknowledgments

During the wheat germplasm collection, we thank for the selfless help from the staff in different research laboratories and experimental stations of the China Agricultural Research System. We also thank the professors and doctors of the Soil Fertility and Plant Nutrition Team of Northwest A&F University, for their critical comments and constructive suggestions.

## 12 Figure Legends

## 13 Supplementary Material

**Supplementary Figure 1**. Relationships of wheat grain Zn concentration to the biomasses of glume (A) and stem (B) and Zn concentrations of glume (C) and stem (D) for the 19 high-yield cultivars over the three experimental years. Determination coefficients (*P*-value) indicate the relationship of grain Zn concentration to each variable.

**Supplementary Table 2**. Tested wheat cultivars in the three-year field experiment, with the identified 19 high-yield ones in red color.

**Supplementary Table 2.**
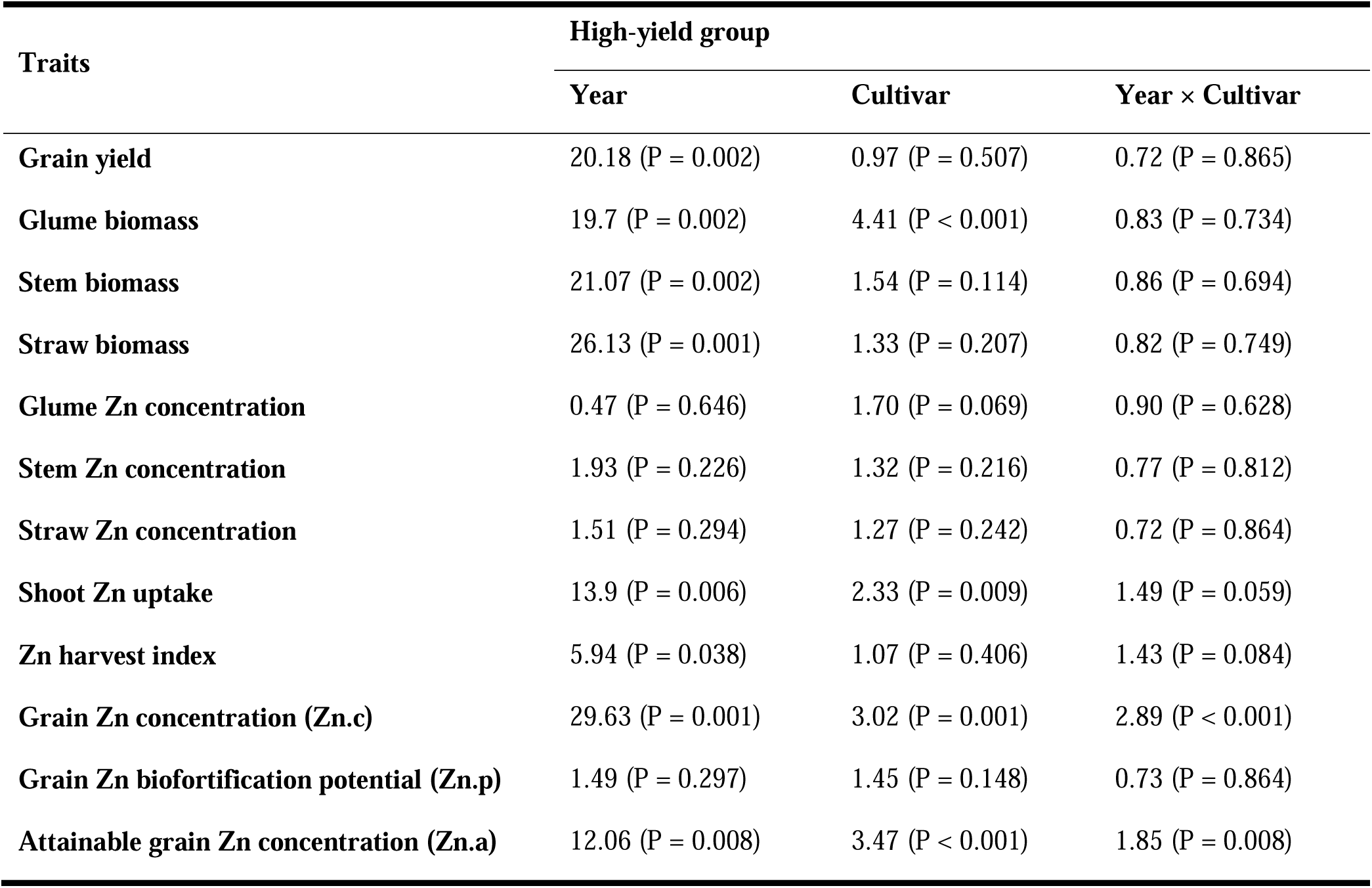
Analysis of variance (ANOVA) for the effects of year, cultivar, and year × cultivar interaction on the different traits of high-yield cultivars.

