## Supplemental Figure 1 and Table 1 for "Quantify the Requirements to Achieve Grain Zn Biofortification of High-yield Wheat on Calcareous Soils"

Supplementary Material

### 1 Supplementary Figures and Tables

#### 1.1 Supplementary Figures

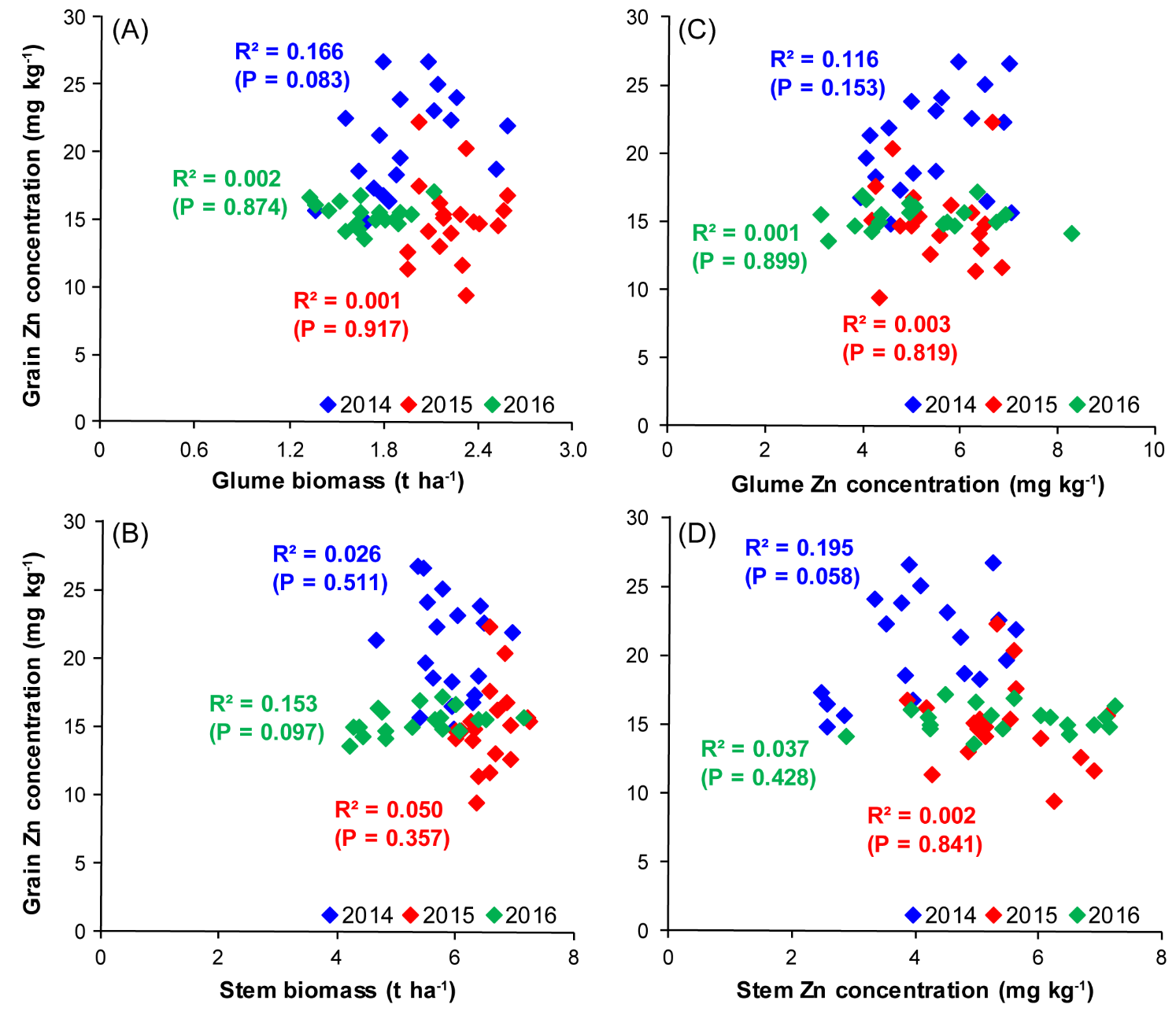

**Supplementary Figure 1.** Relationships of wheat grain Zn concentration to the biomasses of glume (A) and stem (B) and Zn concentrations of glume (C) and stem (D) for the 19 high-yield cultivars over the three experimental years. Determination coefficients (*P*-value) indicate the relationship of grain Zn concentration to each variable.

#### 1.2 Supplementary Tables

**Supplementary Table 1.** Tested wheat cultivars in the three-year field experiment, with the identified 19 high-yield ones in red color.

| **Code** | **Cultivar** | **Released**  **year** | **Code** | **Cultivar** | **Released**  **year** | **Code** | **Cultivar** | **Released**  **year** |
| --- | --- | --- | --- | --- | --- | --- | --- | --- |
| 001 | *Heng 4399* | 2008 | 042 | *Huaimai 25* | 2007 | 083 | *Zhoumai 27* | 2011 |
| 002 | *Hengguan 35* | 2004 | 043 | *Wanmai 68* | 2010 | 084 | *Wenmai 6* | 2000 |
| 003 | *Heng 136* | 2009 | 044 | *Shannong 20* | 2010 | 085 | *Aikang 58* | 2005 |
| 004 | *Gaoyou 2018* | 2008 | 045 | *Huaimai 22* | 2007 | 086 | *Zhoumai 22* | 2007 |
| 005 | *Han 5093* | 2010 | 046 | *Yannong 5158* | 2007 | 087 | *Pingan 5* | 2011 |
| 006 | *Han 6172* | 2001 | 047 | *Yannong 21* | 2002 | 088 | *Baofeng 7228* | 2000 |
| 007 | *Shimai 15* | 2005 | 048 | *Su 553* | 2011 | 089 | *Guomai 0319* | 2009 |
| 008 | *Jimai 32* | 1992 | 049 | *Shiyou 20* | 2009 | 090 | *Hemai 1* | 2006 |
| 009 | *Jimai 22* | 1985 | 050 | *Linmai 4* | 2006 | 091 | *Zhengnong 17* | 2004 |
| 010 | *Liangxing 99* | 2004 | 051 | *Liangxing 66* | 2008 | 092 | *Zhoumai 18* | 2004 |
| 011 | *Jinmai 90* | 2011 | 052 | *Hemai 17* | 2011 | 093 | *Yangfumai 2* | 2003 |
| 012 | *Yaomai 16* | 2011 | 053 | *Liangxing 77* | 2010 | 094 | *Emai 580* | 2012 |
| 013 | *Yunhan 20410* | 2007 | 054 | *Linmai 2* | 2004 | 095 | *Xiangmai 25* | 2008 |
| 014 | *Lin Y8159* | 2013 | 055 | *Shannong 15* | 2006 | 096 | *Emai 596* | 2009 |
| 015 | *Yunhan 618* | 2010 | 056 | *Weimai 8* | 2003 | 097 | *Zhengmai 9023* | 2001 |
| 016 | *Yunhan 719* | 2009 | 057 | *Yannong 0428* | 2008 | 098 | *Chuanmai 104* | 2012 |
| 017 | *Yunhan 805* | 2011 | 058 | *Qingnong 2* | 2010 | 099 | *Xikemai 2* | 2005 |
| 018 | *Yunhan 21-30* | 2003 | 059 | *Yannong 999* | 2011 | 100 | *Neimai 11* | 2007 |
| 019 | *Yunhan 22-33* | 2005 | 060 | *Jimai 22* | 2006 | 101 | *Neimai 836* | 2004 |
| 020 | *Luohan 6* | 2006 | 061 | *Shannong 14* | 2006 | 102 | *Chuanmai 107* | 2000 |
| 021 | *Luohan 7* | 2007 | 062 | *Heima 1* | 2004 | 103 | *Linmai 6* | 2003 |
| 022 | *Luohan 9* | 2009 | 063 | *Luyuan 502* | 2011 | 104 | *Yunmai 56* | 2008 |
| 023 | *Luohan 11* | 2008 | 064 | *Taishan 22* | 2004 | 105 | *Shaan 509* | 2011 |
| 024 | *Luohan 13* | 2009 | 065 | *Jinan 17* | 1999 | 106 | *Zhongmai 895* | 2012 |
| 025 | *Changmai 251* | 2011 | 066 | *Tainong 18* | 2008 | 107 | *Xiaoyan 22* | 2003 |
| 026 | *Changmai 6697* | 2008 | 067 | *Luomai 4* | 2003 | 108 | *Xinong 3517* | 2008 |
| 027 | *Shimai 19* | 2009 | 068 | *Zhoumai 16* | 2002 | 109 | *Xinong 223* | 2012 |
| 028 | *Triticale* | 1980 | 069 | *Zhongyu 10* | 2007 | 110 | *Xiaoyan 15* | 1995 |
| 029 | *Yangmai 13* | 2002 | 070 | *Jimai 20* | 2003 | 111 | *9418* | 2000 |
| 030 | *Ningmai 13* | 2005 | 071 | *Xiangmai 969* | 2006 | 112 | *Shaanmai 139* | 2011 |
| 031 | *Ningmai 14* | 2006 | 072 | *Yumai 58* | 2001 | 113 | *Xinong 9871* | 2008 |
| 032 | *Xinong 979* | 2005 | 073 | *Yumai 18-99* | 2003 | 114 | *Changhan 58* | 2004 |
| 033 | *Huamai 5* | 2010 | 074 | *Yanzhan 4110* | 2003 | 115 | *Changwu 521* | 2008 |
| 034 | *Sumai 188* | 2012 | 075 | *Rumai 0319* | 2009 | 116 | *Chang 134* | 1997 |
| 035 | *Taikong 6* | 2003 | 076 | *Pumai 9* | 2004 | 117 | *Jinmai 47* | 1995 |
| 036 | *Shenmai 22* | 2010 | 077 | *Zhoumai 26* | 2012 | 118 | *Chang 6359* | 2005 |
| 037 | *Xinfumai 1* | 2007 | 078 | *Xuke 1* | 2007 | 119 | *Bei 1* | 2013 |
| 038 | *An 0817* | 2016 | 079 | *Xinmai 26* | 2010 | 120 | *Bei 4* | 2013 |
| 039 | *Longping 203* | 2013 | 080 | *Fanmai 8* | 2008 | 121 | *Bei 9* | 2013 |
| 040 | *Yannong 19* | 2001 | 081 | *Zhoumai 24* | 2009 | 122 | *Xikemai 6* | 2008 |
| 041 | *Xinmai 8* | 2011 | 082 | *Zhengmai 7698* | 2011 | 123 | *Yumai 13* | 2010 |

**Supplementary Table 2.** Analysis of variance (ANOVA) for the effects of year, cultivar, and year × cultivar interaction on the different traits of high-yield cultivars.

| **Traits** | **High-yield group** | | |
| --- | --- | --- | --- |
|  | **Year** | **Cultivar** | **Year × Cultivar** |
| **Grain yield** | 20.18 (P = 0.002) | 0.97 (P = 0.507) | 0.72 (P = 0.865) |
| **Glume biomass** | 19.7 (P = 0.002) | 4.41 (P < 0.001) | 0.83 (P = 0.734) |
| **Stem biomass** | 21.07 (P = 0.002) | 1.54 (P = 0.114) | 0.86 (P = 0.694) |
| **Straw biomass** | 26.13 (P = 0.001) | 1.33 (P = 0.207) | 0.82 (P = 0.749) |
| **Glume Zn concentration** | 0.47 (P = 0.646) | 1.70 (P = 0.069) | 0.90 (P = 0.628) |
| **Stem Zn concentration** | 1.93 (P = 0.226) | 1.32 (P = 0.216) | 0.77 (P = 0.812) |
| **Straw Zn concentration** | 1.51 (P = 0.294) | 1.27 (P = 0.242) | 0.72 (P = 0.864) |
| **Shoot Zn uptake** | 13.9 (P = 0.006) | 2.33 (P = 0.009) | 1.49 (P = 0.059) |
| **Zn harvest index** | 5.94 (P = 0.038) | 1.07 (P = 0.406) | 1.43 (P = 0.084) |
| **Grain Zn concentration (Zn.c)** | 29.63 (P = 0.001) | 3.02 (P = 0.001) | 2.89 (P < 0.001) |
| **Grain Zn biofortification potential (Zn.p)** | 1.49 (P = 0.297) | 1.45 (P = 0.148) | 0.73 (P = 0.864) |
| **Attainable grain Zn concentration (Zn.a)** | 12.06 (P = 0.008) | 3.47 (P < 0.001) | 1.85 (P = 0.008) |

Data in table are ANOVA F values (Pr > F).
